# Methanol fixation is the method of choice for droplet-based single-cell transcriptomics of neural cells

**DOI:** 10.1101/2022.08.03.502652

**Authors:** Ana Gutiérrez-Franco, Mohamed N. Hassan, Loris Mularoni, Mireya Plass

## Abstract

Single-cell transcriptomics methods have become very popular to study the cellular composition of organs and tissues and characterize the expression profiles of the individual cells that compose them. The main critical step in single-cell transcriptomics is sample preparation. Several methods have been developed to preserve cells after sample dissociation to uncouple sample handling from library preparation. Yet, the suitability of these methods depends on the types of cells to be processed. In this project, we perform a systematic comparison of preservation methods for droplet-based single-cell RNA-seq (scRNA-seq) on neural and glial cells derived from induced pluripotent stem cells (iPSCs) and highlight their strengths and weaknesses. We compared the cellular composition and expression profile of single-cell suspensions from fresh NPCs with that of NPCs preserved with Dimethyl Sulfoxide (DMSO), Methanol, vivoPHIX and Acetil-methanol (ACME). Our results show that while DMSO provides the highest cell quality in terms of RNA molecules and genes detected per cell, it strongly affects the cellular composition and the expression profile of the resulting datasets. In contrast, methanol fixed samples display a cellular composition like that of fresh samples while providing a good cell quality and smaller expression biases. Taken together, our results show that methanol fixation is the method of choice for performing droplet-based single-cell transcriptomics experiments on neural cell populations.

## Introduction

Single-cell transcriptomics (scRNA-seq) methods have revolutionized the way we study in a high-throughput manner the expression of genes across individuals, tissues and even in disease^1–3^. Previously, studies were limited to identify changes in the expression levels of genes in bulk, that is, in the population of cells that composed a particular sample. Thus, these approaches mixed two different effects: changes in the cell composition of the sample of interest and changes in the expression of genes within individual cells. Now, scRNA-seq allows assessing these two effects independently and can detect both changes in the cellular composition^4,5^ and the expression of genes in specific cell types^6,7^.

Despite the popularity of scRNA-seq methods, there are still several technical challenges unsolved. For instance, the dissociation of the cells from a tissue and the obtention of a good cell suspension, necessary for scRNA-seq, is highly tissue specific and may require the use of different strategies including enzymatic digestion, mechanical disgregation, fluorescence-activated cell sorting (FACS), and other technologies^8–12^. As a result, the preparation of samples for scRNA-seq can take several hours and makes it difficult to couple sample preparation with the acquisition of the transcriptomes of thousands of cells.

Several methods have already been developed to overcome this problem and uncouple sample handling from library preparation. Among these protocols, we find both homemade and commercial solutions including methanol fixation^13,14^, DSP^15^, DMSO^16,17^ cryopreservation^17^, ACME^10^, PFA^18^, CellCover^17^ and vivoPHIX^19^. However, many of these protocols have been tested only in cell lines or easy-to-obtain cells such as peripheral blood cells (PBMCs) and thus, it is not clear how its performance is in cells that are difficult to dissociate or that are potentially damaged during dissociation process such as neural cells.

In this work, we have compared the performance of five popular fixation and preservation methods in neural and glial cells derived from human induced pluripotent stem cells (hiPSCs). The results from our work show that the different preservation/fixation methods affect the samples in different ways, including biases in the transcriptomic profile, cell composition and library complexity. DMSO cryopreservation provides the highest cell quality in terms of library complexity. Yet, the obtained datasets are strongly depleted of neurons and display a stronger stress signature. In contrast, ACME and vivoPHIX do not significantly affect the cell composition of the single-cell suspensions, but damage the RNA, which reduces the library complexity and thus the number of genes and RNA molecules detected in individual cells. Taken together, our results show that methanol fixation is the method of choice for performing droplet-based single-cell transcriptomics experiments on neural cell populations as it provides a high library complexity without affecting the cell composition nor gene expression in comparison to fresh samples.

## Results

### Experimental set up for the systematic comparison of preservation protocols

Individual or pooled iPSC cell lines were differentiated to cortical neurons using a previously described protocol^20^ with minor modifications. Briefly, hiPSC colonies were seeded in 12 well-plates coated with matrigel and after reaching confluency, neural differentiation was induced using a combination of BMP inhibitors (noggin, dorsomorphin and SB431542) (Figure 1a). After 29 - 50 days of differentiation, cells were dissociated with papain-accutase solution to obtain a single-cell suspension. At this point, some of the samples were directly encapsulated using Dolomite Bio automatic Drop-seq set-up NADIA (Fresh), cryopreserved with DMSO^17^ or preserved using different chemical compounds that stabilize RNA molecules, some of which have been previously used in single-cell transcriptomics including Methanol^14,21^, ACME^10^, vivoPHIX^19^ and Cell Cover^17^ (Figure 1b).

**Figure 1.**
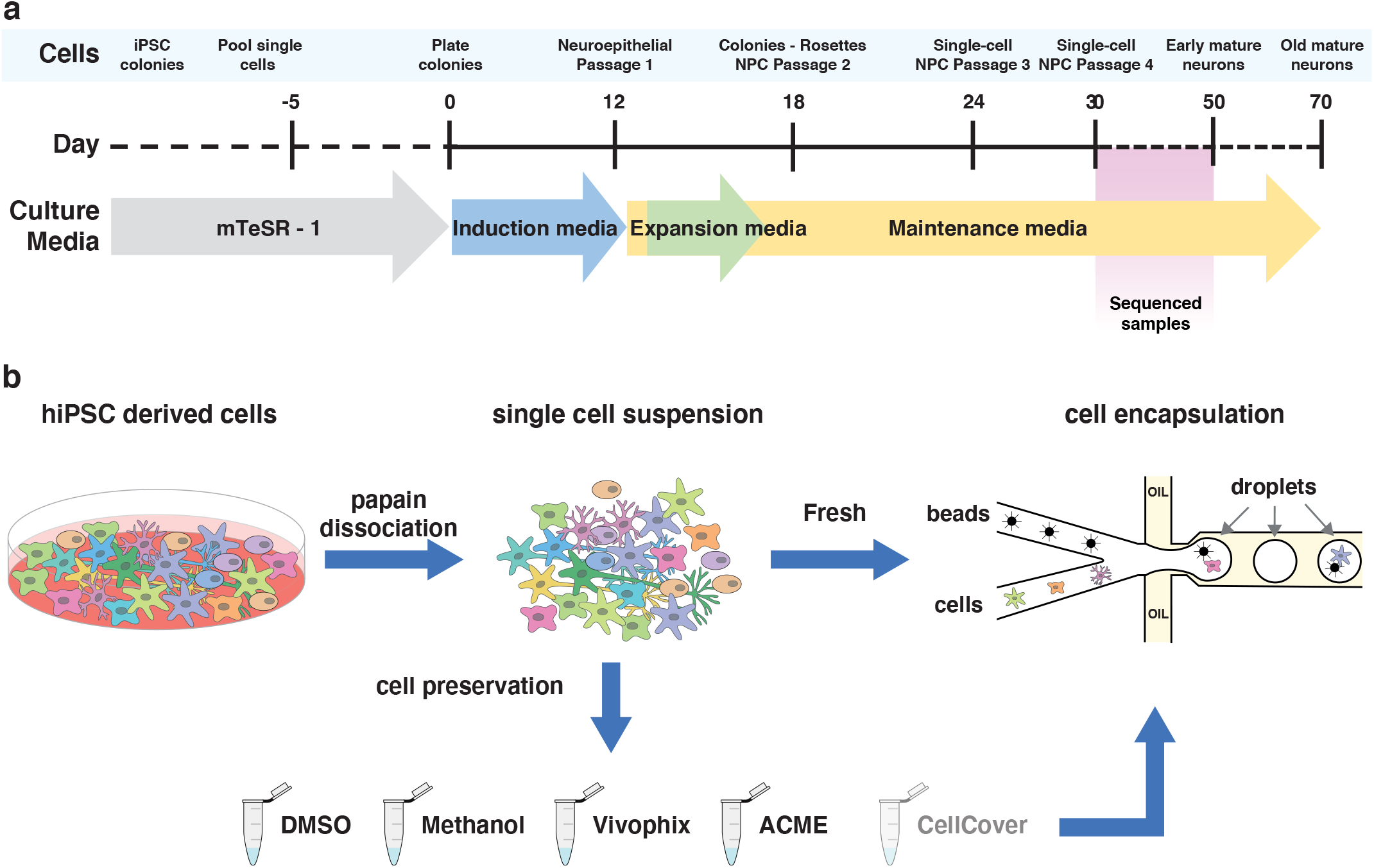
Experimental set up. **a)** Schematic representation of the differentiation of hiPSCs to neural progenitor cells. **b)** Systematic comparison of preservation methods. After differentiation of hiPSCs, all cells were dissociated using papain-accutase and either directly encapsulated or preserved using one of the different reagents tested. After being preserved, cells were thawed or rehydrated and encapsulated using a commercial Drop-seq set up using the same protocols.

### Cell preservation methods can impact RNA quality

Before performing single-cell transcriptomics analysis, we investigated if preservation of single-cell suspensions affects the quality of obtained RNA. For that purpose, we extracted total RNA from fresh and preserved NPCs stored at -80ºC or 4ºC for up to 15 days (Table S1). For each of the samples, we quantified the total amount of RNA and assessed its quality using the Agilent TapeStation system. Our results showed that the quality of the RNA extracted depended on the preservation method used. DMSO, Methanol and ACME samples had very high RIN values (∼9), similar to those of Fresh samples. In contrast, the vivoPHIX sample had some RNA degradation (RIN ∼7) and the samples processed with CellCover had stronger degradation levels, with RIN values ∼2 at 4ºC and ∼6 at -80ºC (Figure S1). These results demonstrate that CellCover is not suitable for long-term storage of cells for single-cell transcriptomics. Taking this into consideration, we decided to discard CellCover for the systematic comparison of preservation methods.

### Cell preservation affects the complexity of captured single-cell transcriptomes

To compare the impact of different preservation methods on neural progenitor cell (NPC) populations, hiPSCs were differentiated to NPCs as previously described and either preserved using one of the different protocols or directly encapsulated with the Nadia equipment (Figure 1b). Cell encapsulation and library preparation was done following the same protocol for all samples. Libraries were sequenced using Nextseq 550 sequencing platform and processed using Drop-seq tools^22^ to obtain a Digital Expression Matrix (DGE) for each of them. We evaluated the quality of the libraries obtained using different metrics to assess library complexity, cell damage and the overall quality of the samples.

We initially examined the quality of the libraries by looking at the trace of the cDNA obtained for each sample. Manual inspection of the profiles showed that ACME and vivoPHIX samples had less cDNA and smaller fragments (average fragment size 1187 and 1079 respectively) than the rest of the libraries (Figure 2a), which is consistent with RNA degradation. This was expected in the vivoPHIX samples, which showed already a lower RNA quality after being preserved for 2 weeks, but not for ACME samples, which had RIN values similar to that of Fresh samples (Figure S1). Next, we evaluated if RNA degradation impacted the complexity of the libraries. We used *samtools*^23^ to downsample each of the unaligned BAM files to produce files containing 10%, 20%, 30%, etc. of the original dataset. Each of these subsamples was then processed using the same computational pipeline used to generate DGEs. Our results show that at the same sequencing depth, DMSO cryopreserved cells and methanol fixed cells yield a comparable or higher number of Genes and UMI per cell than Fresh cells. In comparison, at 7500 reads per cell, ACME and vivoPHIX samples have around 40% of the Genes and UMIs obtained in fresh cells (Figure 2b) (Table 2). The higher number of genes and UMIs in the DMSO and M3 and M4 methanol fixed samples is due to differences in the capture efficiency of the beads used for the encapsulation and not to an intrinsic higher capture efficiency in these conditions (Figure S2 and Table S1).

**Figure 2.**
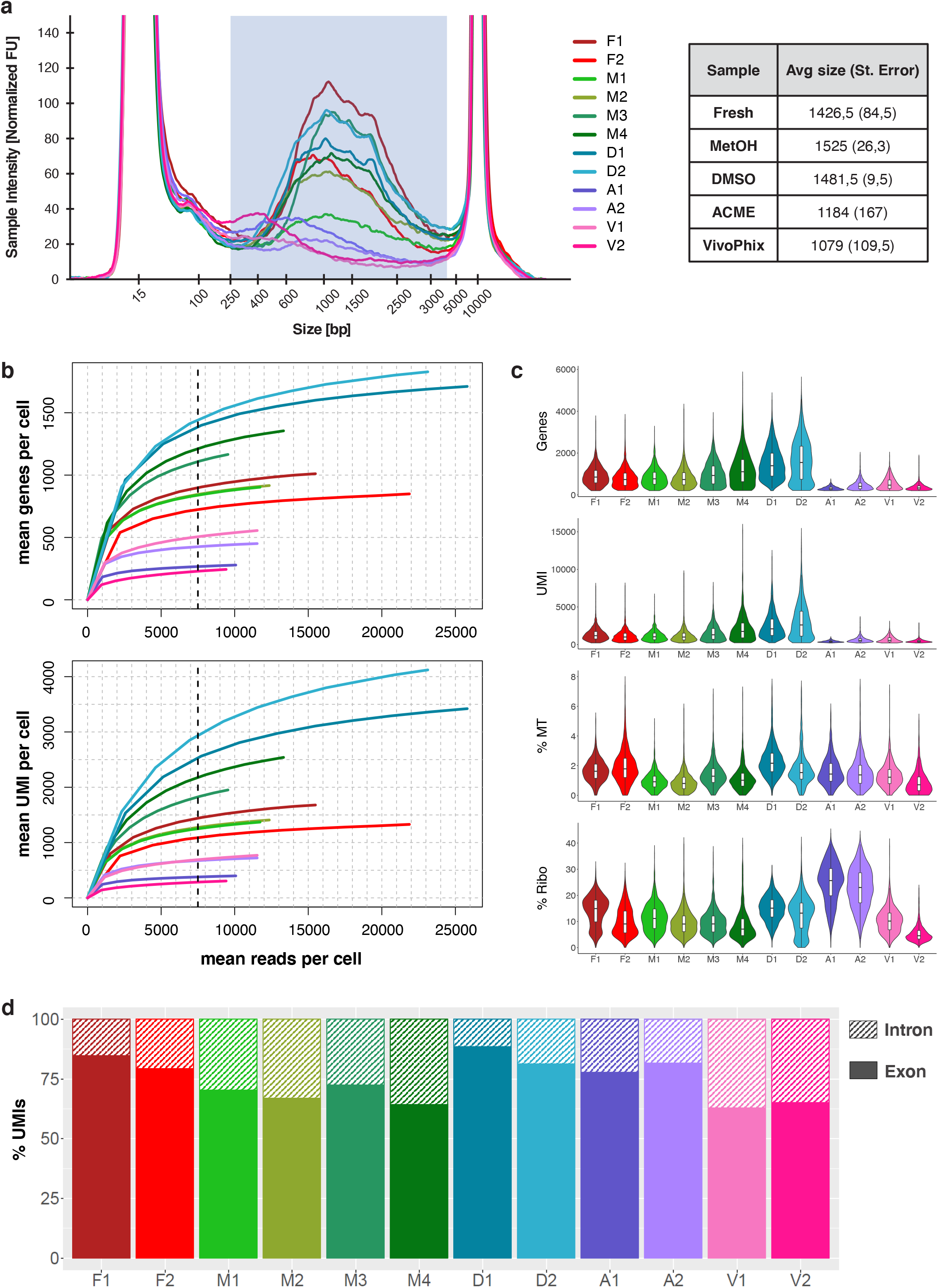
Systematic comparison of the effects of cellular preservation on library quality. **a)** Trace of the libraries obtained after encapsulation including Fresh Samples (F1 and F2), methanol fixed samples (M1, M2, M3 and M4), DMSO cryopreserved samples (D1 and D2), ACME fixed samples (A1 and A2) and vivoPHIX samples (V1 and V2). The libraries of the samples fixed with vivoPHIX and ACME have smaller fragment sizes and less RNA, which is consistent with RNA degradation. **b)** Downsampling plots showing the number of genes (top) and UMIs (bottom) *as a function of sequencing depth (mean reads per cell)*. In both cases, DMSO (blue) and Methanol (green) samples have a depth equivalent or superior as that of fresh samples (red). **c)** Violin plots showing the distribution in the amount of Genes, UMIs, mitochondrial content (% MT) and ribosomal content of the final filtered cells. **d)** Barplots showing the percentage of intronic and exonic reads of each of the libraries.

The lower complexity of vivoPHIX and ACME libraries is also reflected in the proportion of low quality cells in the samples, which have few UMIs and Genes detected per cell (Figure 2c, Table 1), and may correspond to empty droplets or droplets containing broken cells^24^. While the number of low-quality cells discarded is around 10% of cells in Fresh, DMSO and Methanol fixed cells, this number increases to up to 30% and 40% in ACME and vivoPHIX samples respectively (Table 1).

**Table 1.** Cell quality statistics.

| Sample | Cells | Filtered Cells | % cells kept | Mean genes per cell | Mean UMI per cell | % MT Genes | % Ribo Genes |
| --- | --- | --- | --- | --- | --- | --- | --- |
| F1 | 1632 | 1565 | 95,89 | 908 | 1331.36 | 1.65 | 14.03 |
| F2 | 1227 | 1098 | 89,49 | 793 | 1145.34 | 1.93 | 10.08 |
| M1 | 1703 | 1556 | 91,37 | 834 | 1167.05 | 0.97 | 11.40 |
| M2 | 1621 | 1521 | 93,83 | 818 | 1160.68 | 0.89 | 9.55 |
| M3 | 1696 | 1660 | 97,88 | 1001 | 1521.20 | 1.40 | 9.59 |
| M4 | 1207 | 1139 | 94,37 | 1239 | 2149.08 | 1.16 | 8.66 |
| D1 | 1141 | 1129 | 98,95 | 1512 | 2513.08 | 2.32 | 14.93 |
| D2 | 1241 | 1222 | 98,47 | 1625 | 3085.38 | 1.78 | 12.56 |
| A1 | 1215 | 545 | 44,86 | 315 | 403.84 | 1.62 | 24.82 |
| A2 | 1410 | 1014 | 71,91 | 460 | 637.01 | 1.53 | 22.76 |
| V1 | 1455 | 1013 | 69,62 | 537 | 674.95 | 1.32 | 10.42 |
| V2 | 2824 | 1126 | 39,87 | 364 | 452.94 | 0.89 | 5.31 |

**Table 2.** Summary of the effects of preservation methods on single-cell libraries.

| Fixation Method | Library yield | Library complexity | Low quality cells | Composition bias | Stress signature | Expression Bias |
| --- | --- | --- | --- | --- | --- | --- |
| DMSO | HIGH | HIGH | FEW | STRONG | HIGH | LOW |
| Methanol | HIGH | HIGH | FEW | WEAK | LOW | LOW |
| ACME | LOW | LOW | MANY | WEAK | LOW | MEDIUM |
| VivoPHIX | LOW | LOW | MANY | WEAK | LOW | MEDIUM |

After discarding low quality cells, the remaining vivoPHIX and ACME cells still show clear differences in quality (Figure 2c, Table 1). Interestingly, the % of UMIs mapped to mitochondrial genes was similarly low across all samples, suggesting that the lower number of captured RNAs in these samples may not be due to RNA leakage or cellular damage^24^ but rather be related with the capture efficiency of the RNAs itself or RNA degradation. Additionally, ACME samples showed a much higher fraction of UMIs mapped ribosomal proteins than any of the other samples (Figure 2c), which has also been previously associated to low quality cells or technical artifacts^25^.

One of the reasons why vivoPHIX and ACME fixed cells have less RNAs per cell could be that the fixation method facilitates cell breakage. In this case, we would expect lower library complexity and a higher fraction of reads coming from intronic regions, which mainly come from pre-mRNAs and are enriched in nuclei^26^. In our samples, the fraction of intronic reads coming from each sample was variable across preservation methods, ranging from 14% in DMSO samples to almost 40% in vivoPHIX (Figure 2d). Thus, the high fraction of intronic reads in vivoPHIX samples can indicate a higher fraction of broken cells, which can partially explain the lower library complexity of these samples.

### DMSO cryopreservation alters the cell composition of iPSC derived cell populations

After assessing the overall quality metrics of the samples, we investigated whether preservation methods affect the cellular composition of the samples. For that purpose, we pooled all the samples and analyzed them together. Initial analysis showed a strong batch effect driven mainly by the preservation method used (Figure S3). Thus, we used Harmony^27^ to integrate the datasets and identify the cell populations obtained from hiPSC differentiation (Figure 3). After batch correction, we identified 13 cell populations corresponding to proliferating progenitors, NPCs, astroglial precursors, intermediate progenitors and different types of neurons characterized by the expression of specific marker genes (Figure 3, Figure S4, S5) (Table S4, S5). All the clusters identified were present in all the individual samples. Yet, the relative abundance of each of the cell populations changed depending on the preservation method used (Figure 4a). To investigate if these changes were due to experimental biases or they were systematic biases due to the fixation method, we performed a compositional analysis of the samples with scCODA^5^. The results from this analysis highlight a significant depletion of excitatory neurons and an enrichment in astroglial progenitors in DMSO cryopreserved samples compared with Fresh samples while we do not find significant compositional biases in the samples preserved using any of the other methods (Figure 4b).

**Figure 3.**
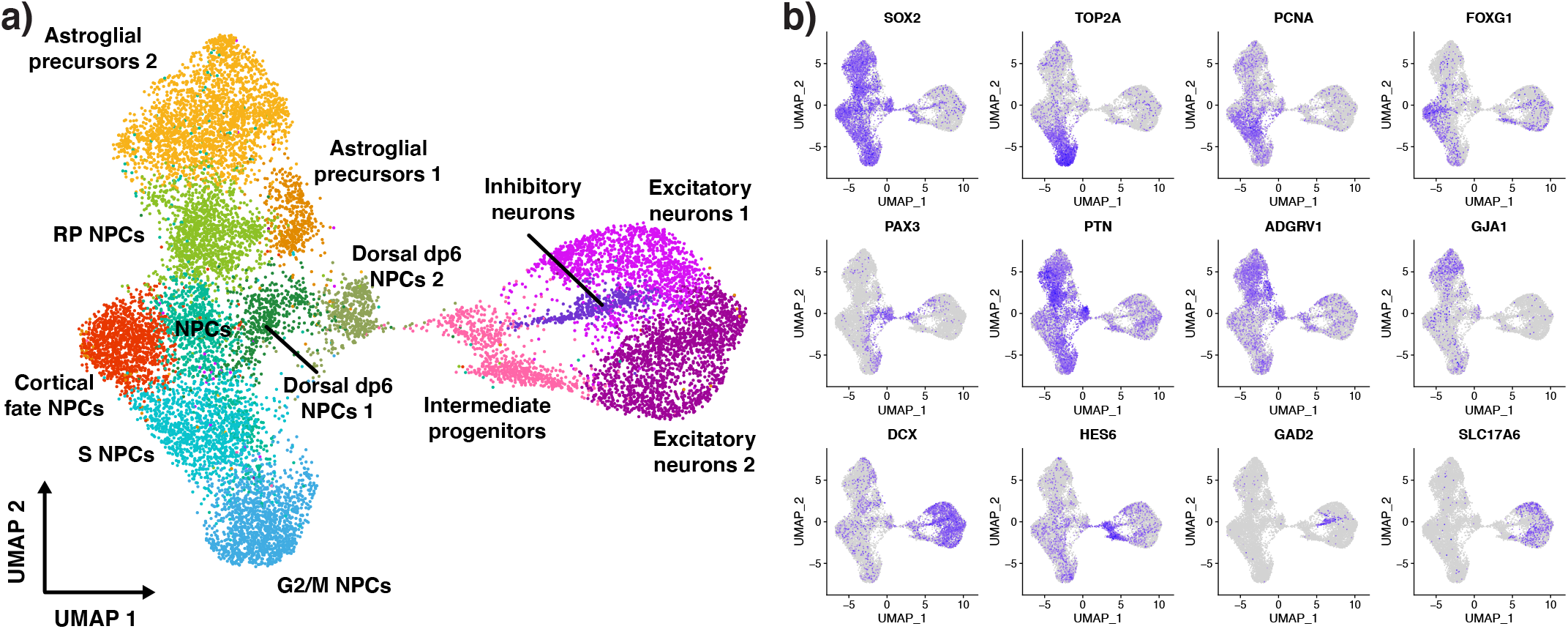
scRNA-seq identifies multiple diverse neural and astroglial progenitors and immature neuron populations. **a)** UMAP plot showing the cell populations identified in the NPC samples, which include different types of NPC populations, astroglial precursors, intermediate progenitors and both excitatory and inhibitory immature neurons. **b)** Feature plots of known markers that have been used to identify the cell populations in **a** including NPC markers (SOX2), proliferation markers (TOP2A and PCNA), cortical fate markers (FOXG1), dorsal fate markers (PAX3), roof plate markers (PTN), astroglial markers (ADGRV1 and GJA1), immature neuronal markers (DCX), intermediate progenitor markers (HES6), inhibitory (GAD2) and excitatory (SLC17A6) neuronal markers.

**Figure 4.**
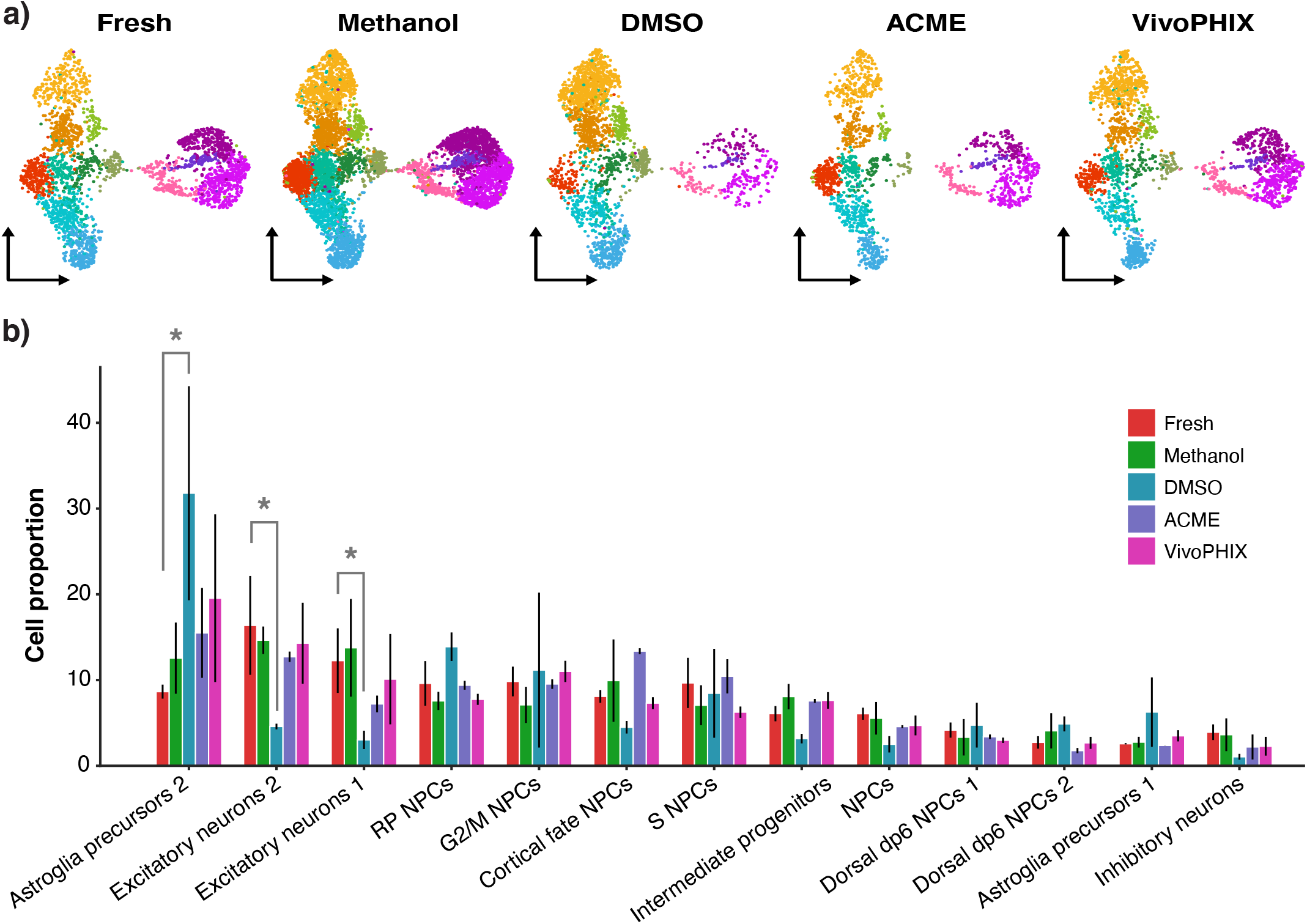
DMSO cryopreservation impacts the cell composition of scRNA-seq datasets. **a)** UMAP plots showing the distribution of cells across clusters for each of the fixation methods. DMSO preserved cells show a strong depletion of neuronal clusters compared to the other methods. Clusters are colored as in Figure 3a. **b)** Barplot showing the proportion of cells assigned to each cluster for each method. The high of the bars represents the average for all samples preserved using the same method while bars represent the standard deviation. * Indicates samples with a significant difference in cell proportion relative to fresh samples as predicted by scCODA.

### Preservation protocol affects gene expression across cell populations

Finally, we investigated if preservation methods induced changes in gene expression. Previous studies have shown that dissociation and preservation methods can induce cellular stress that is reflected at the transcriptomic level^9,28^. Accordingly, we checked the expression of Immediate Early genes (IEGs) and apoptosis markers in our datasets. DMSO cryopreserved cells showed higher expression of IEGs than all the other samples (Figure 5A), indicating that freezing and thawing stresses cells in a way that is reflected on the transcriptomic profile. In contrast, the apoptosis gene signature was higher in ACME and vivoPHIX samples (Figure 5B). These gene expression changes were minor, as overall gene expression across all samples strongly correlated (Figure S6).

**Figure 5.**
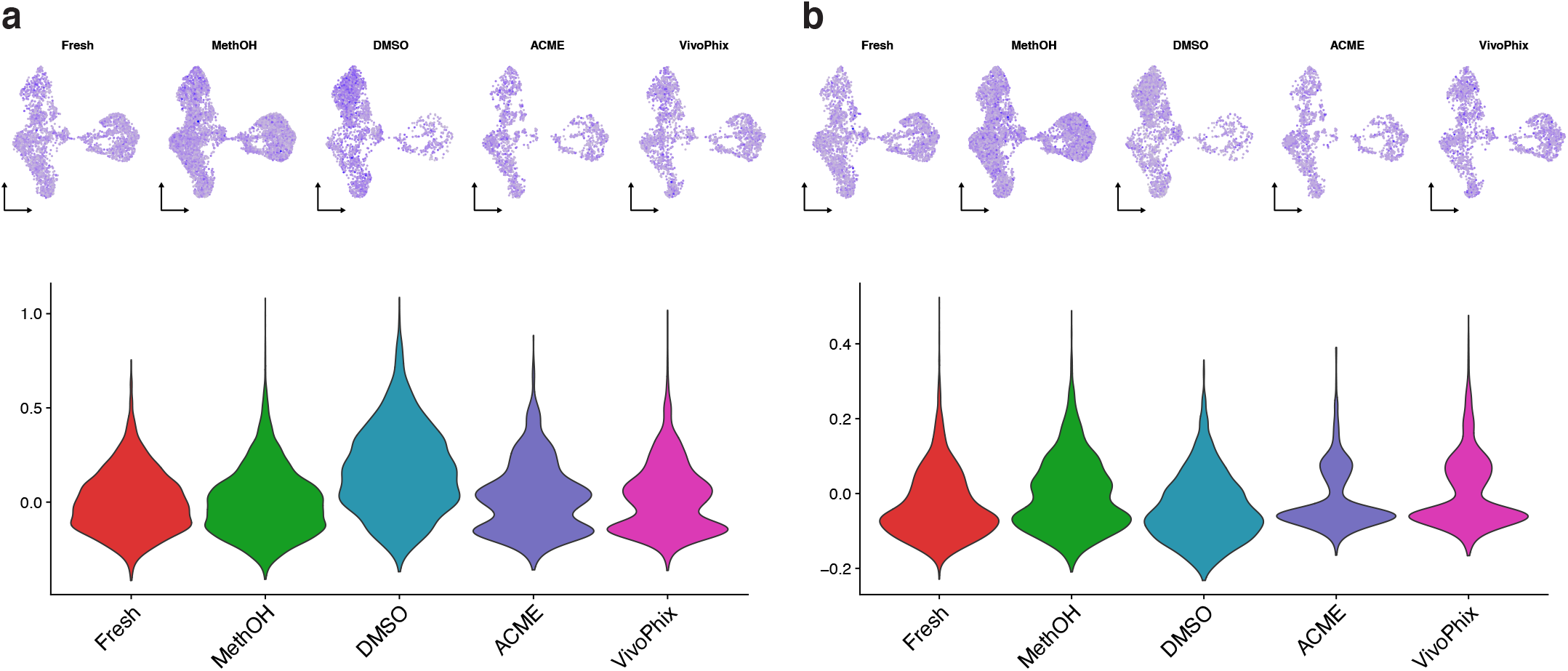
DMSO cryopreservation affects the overall expression of stress markers in scRNA-seq data. **a**,**b)** Feature plots (top) and violin plots (bottom) showing the distribution of stress **(a)** and apoptosis **(b)** gene expression signatures across cells. Both signatures show similar expression across all datasets except for DMSO, which shows higher stress signature across all clusters.

Next, we checked if preservation was driving further gene expression changes that could impact sample comparison. We generated pseudobulk counts for each of the samples and performed a correlation analysis at the cluster level. Our analyses show that the fixation method induces biases in the clustering of cell populations across samples. While in general we observe that cell clusters from the same or similar cell types from methanol, DMSO and fresh sample cluster together, ACME and vivoPHIX cells cluster separately (Figure 6). These analyses demonstrate that methanol fixation impacts less gene expression in comparison to other fixation/preservation methods. Methanol fixed cells show overall a low level of IEGs and apoptosis genes and their transcriptomic profile is close to that of fresh cells.

**Figure 6.**
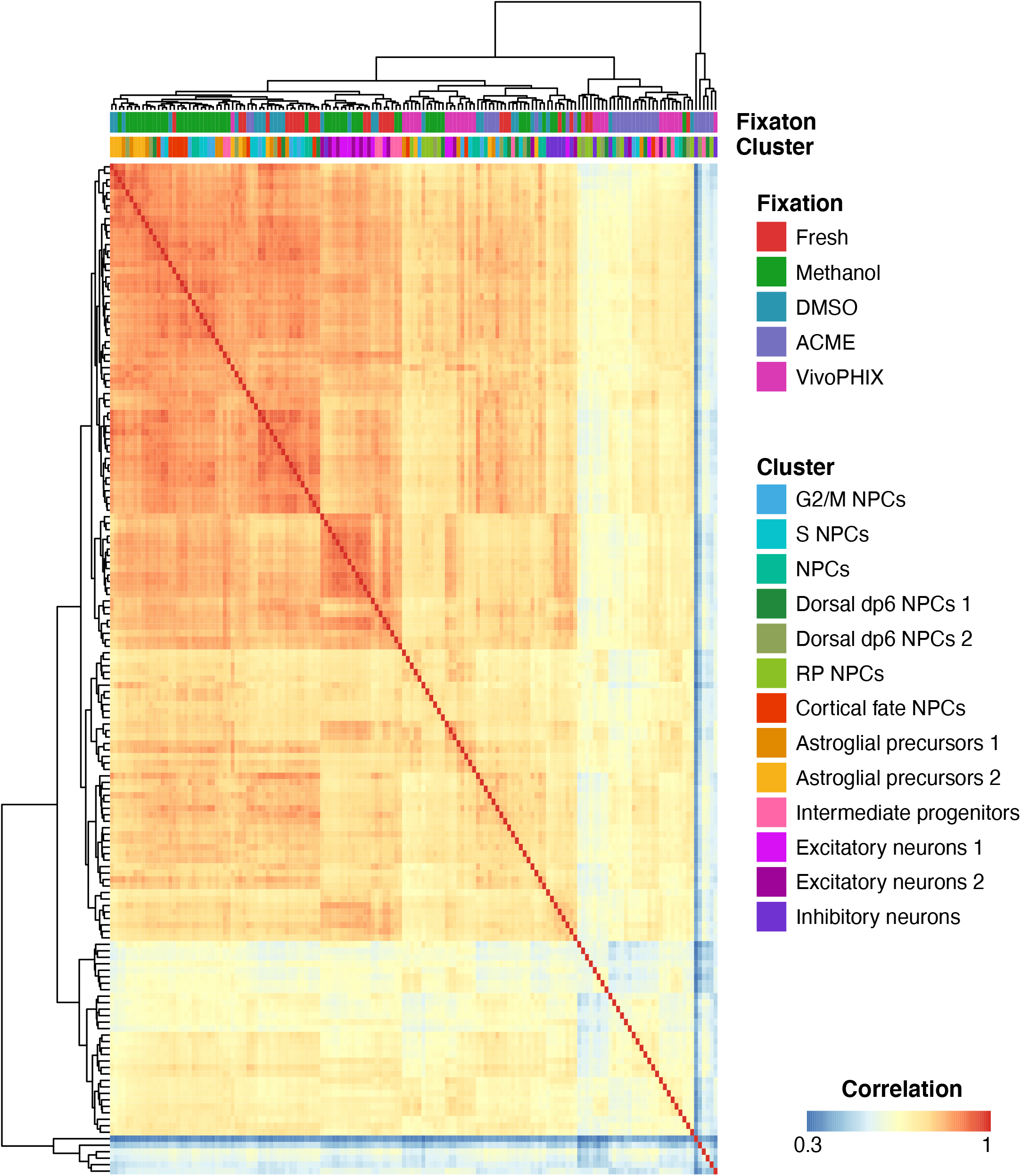
Cluster level expression correlation separates vivoPHIX and ACME clusters. Heatmat plot showing per cluster gene expression correlation across samples. Fixation method and cluster identity for each sample are indicated as color boxes on top of the heatmap. While neuronal and progenitor cell clusters show a good correlation among DMSO, Methanol and Fresh samples, this correlation is lower with vivoPHIX and ACME samples.

## Discussion and Conclusions

Single-cell transcriptomics methods are becoming the new standard to study transcriptomic changes across samples and conditions. These technologies are relatively new compared to bulk transcriptomics methods such as RNA-seq or 3’ seq. Thus, in many cases there are not yet standard preparation protocols for the different samples used. In this project, we have compared how commonly used preservation and fixation methods affect the cell composition and expression of neural and glial cell populations derived from hiPSCs. This work thus extends previous studies that have compared the effects of only one preservation method on the quality of single-cell transcriptomes or that have focused on their effects on different cell types and thus may not be applicable to neural cells^13–19^. Our results show that different preservation/fixation methods affect the quality of the single-cell transcriptomics datasets in different ways, including decreased library complexity, changes in cell composition, and alterations in the expression profile of individual cells (Table 2).

In terms of library complexity, ACME and vivoPHIX samples show a strong decrease in the amount of cDNA obtained after single-cell encapsulation (Figure 2a), which is also reflected in a lower detection of Genes and UMIs (Figure 2b). This decrease is consistent with previous reports that show that both ACME and vivoPHIX preserved samples have lower RNA quality than that of fresh cells^10,29^. Additionally, these methods are damaging the cells and/or favoring RNA leakage, which is reflected in a higher fraction of ribosomal reads (ACME samples) and intronic reads (vivoPHIX). Additionally, this lower cell complexity due to a higher dropout rate is likely the cause of the biases in the cell population clustering analysis (Figure 6).

When looking at cell composition, our results clearly highlight a strong depletion of neuronal cells in the DMSO cryopreserved samples (Figure 4). While DMSO has been previously reported as an excellent method of cell preservation for single-cell transcriptomics^17,30^, none of these studies looked at the effect of DMSO on mature neuronal cells. The reduction in the number of recovered neurons is probably due to DMSO toxicity^31,32^ or DMSO-induced reactive gliosis, which has been reported previously and can affect cells after a very brief exposure^31^. Yet, the obtained neurons do not show strong expression biases as they cluster with the same populations obtained from fresh and methanol fixed samples (Figure 6). While our results demonstrate that DMSO is not a good choice for performing compositional analysis of iPSC-derived cells, it could be used for profiling pure neural samples when cell availability is not an issue.

Taking into account the pros and cons of the different methods (Table 2), our comparative analysis indicates that methanol fixation is the best method of preservation to perform single-cell transcriptomics analyses on neural cells. Libraries from methanol fixed cells have similar complexity to that of fresh cells and do no present strong biases in gene expression (Figure 6) or cell composition (Figure 4), thus providing the sample the most similar results to that of Fresh cells.

## Methods

### iPSC cell culture and differentiation

Human iPSCs were maintained on 1:40 matrigel (Corning, #354277) coated dishes in supplemented mTeSR-1 medium (StemCell Technologies, #85850) with 500 U/ml peniciline and 500 mg/ml streptomycin (Gibco, #15140122). For the differentiation of cortical neurons the protocol described previously^20^ was followed with slight modifications. Briefly, hiPSC colonies were seeded in 12 well-plates coated with 1:40 matrigel at a cell density sufficient to ensure 100% confluence one day after plating. At day 1, the medium was switched to neural induction medium (neural maintenance medium (1:1 ratio of DMEM/F-12 GlutaMAX (Gibco, #10565018) and Neurobasal (Gibco, #21103049) medium with 1x N-2 (Gibco, #17502048), 1x B-27 (Gibco, #17504044), 5 μg/ml insulin (Sigma, #I9278), 1 mM L-glutamine (Gibco, #35050061), 100 μM non-essential amino acids (Lonza, #BE13-114E), 100 μM 2-mercaptoethanol (Gibco, #31350010), 50 U/ml penicillin and 50 mg/ml streptomycin) supplemented with 500 ng/ml noggin (R&D Systems, # 3344-NG-050), 1 μM Dorsomorphin (StemCell technologies, #72102) and 10 μM SB431542 (Calbiochem, # 616461)). The neural induction media was replaced every day for 9-12 days until the neuroepithelial sheet was formed. At this point, the neuroepithelial cells were collected in aggregates using dispase (StemCell Technologies, #07923) and seeded on 20 μg/ml laminin-coated (Sigma, #L2020) 6 well-plate containing 2 ml of neural maintenance medium. Cells were incubated in neural maintenance medium with every-other day replacement until neural rosette structures were recognizable (days 12-15 after neural induction). Then, 20 ng/ml of bFGF (Peprotech, #100-18B) was added to the medium for 2-4 days to promote the expansion of neuro stem cells (NSC). At day 18 after neural induction, cells were splitted with dispase for precursors amplification. At day 24, when neurons begin to accumulate at the outside of the rosettes, cells were passaged 1:3 using Accutase (Merck Millipore, #SCR005) in a single-cell suspension and seeded at 50000 cells/cm^2^ on 20 μg/ml laminin-coated 6-well plate. After a week, cells were splitted again (ratio 1:4) and seeded on 20 μg/ml laminin-coated 6-well plates and continued the culture for up to 50 days after neural induction with medium changes every second day. The use of hiPSCs in this work was approved by the Spanish’ National commission of guarantees for the donation and use of human cells and tissues Institute of Health Carlos III.

### Cell dissociation

Cells were dissociated into a single-cell suspension following a previously described protocol optimized for scRNA-seq techniques^33^. In summary, cells were enzymatically dissociated for 35 min at 37ºC using papain-accutase dissociation buffer (1:1) (PDS Kit, Papain, Worthingron Biochemical Corporation, #LK003176), and quenched with DMEM/F-12 GlutaMAX supplemented with 10 µM of ROCK inhibitor (Y-27632, StemCell Technologies, #72304) and 0.033 mg/ml of DNase (DNase (D2), Worthington Biochemical Corporation, #LK003170). Cell suspension was filtered through a 40 µm strainer (Pluriselect Life Science, #43-10040-60) and then centrifuged at 150 g for 3 min at RT. After 3 washes with 0.4 mg/ml BSA in DPBS, cells were counted, and viability by trypan blue method recorded. Only samples with cell viability higher than 80% were included in the study.

### DMSO cryopreserved sample preparation

Around 2.5×10^6^ cells after dissociation were cryopreserved in cryovials in freezing medium (neural maintenance medium supplemented with 10% v/v of dimethyl sulfoxide (DMSO) (Sigma-Aldrich, #D2438) and 20 ng/ml bFGF). The cryovials were placed into a Mr. Frosty freezing container (Nalgene, #5100-001) previously filled up with isopropyl alcohol and stored at -80ºC overnight (ON) and then transferred for long storage to a vapor phase nitrogen freezer. DMSO cryopreserved samples were thawed in a water bath at 37ºC in continuous agitation, then 1 ml of maintenance medium was added to the vial and transferred to a falcon tube with 10 ml of maintenance medium. Cells were centrifuged at 160 g for 5 min at RT. The supernatant was carefully removed, and the cell pellet was washed with 1 ml of DPBS and 0.01% BSA and then transferred to a 1.5 ml DNA LoBind tube (Eppendorf, #022431021). Cells were pelleted again and resuspended in DPBS and 0.01% BSA. Finally, cells were filtered through a 40 µm strainer and counted in a Neubauer chamber using the standard trypan blue method.

### Methanol sample preparation

Following the methanol fixation protocol for single-cell RNA-seq in 10X Genomics (CG000136), ice-cold DPBS was added to resuspend a 2.5×10^6^ cell pellet. Pre-chilled 100 % methanol was added dropwise until the final methanol concentration reached 80%. Samples in DNA LoBind tubes were placed on ice 30 min, then ON at -20ºC and finally transferred to -80ºC for long storage. Methanol fixed cells were thawed on ice and centrifuged to remove the supernatant. The cell pellet was washed and rehydrated in DPBS with 0.01% BSA and 0.2 U/µl of RNase inhibitor (Takara Bio, #2313A) and 1 mM DTT (Sigma-Aldrich, #D0632)) to avoid RNA degradation. Cells were filtered again with a 40 µm strainer and counted in a Neubauer chamber.

### ACME sample preparation

A pellet of 1×10^6^ to 5×10^6^ cells was gently resuspended with 100 µl of wash buffer (DPBS with 0.01% BSA and 0.2 U/µlof RNase inhibitor and 1 mM DTT). Then, ACME solution (wash buffer: methanol: acetic acid: glycerol; in a final ratio of 13:3:2:2) was added dropwise while mixing the tube to a final volume of 1 ml and incubated for 30 min at RT. After centrifugation at 1000 g at 4ºC for 5 min and discarding the supernatant, the fixed cell pellet was washed twice in wash buffer and resuspended in wash buffer supplemented with 10% v/v DMSO for storage at -80ºC. ACME fixed samples were thawed and rehydrated following the same protocol as methanol fixed samples.

### vivoPHIX sample preparation

A cell pellet containing 1×10^6^ to 5×10^6^ cells was gently resuspended with 25 µl of DPBS + 0.01% BSA. For sample fixation, 75 µl of vivoPHIX reagent (Rapid Labs, #RD-VIVO-5) (3:1 ratio) was added and mixed by inverting the tube 10 times during the first 5 min of fixation. Next, the tube was placed on a wheel device at RT and incubated for 30 min more. Samples were stored at 4ºC ON and then transferred to -80ºC for long storage.

vivoPHIX fixed samples were rehydrated by the addition of one volume (100 µl) of 100% ethanol and mixing the tube several times by inversion. Then, cells were pelleted at 1000 g for 5 min at RT and the supernatant was discarded. 0.5 ml of vivoPHIX-SCAA (1 volume of vivoPHIX with 3 volumes of glacial acetic acid) was added very slowly to the cell pellet without disturbing it and incubated for exactly 3 minutes at RT. The vivoPHIX-SCAA from the pellet was removed and cells were pelleted again at 100 g for 5 min at RT to remove any remaining liquid using a P20 pipette tip. The cell pellet was washed three times with DPBS with 0.01% BSA and 0.2 U/µlof RNase inhibitor and 1 mM DTT and the supernatant discarded. Cells were filtered with a 40 µm strainer and counted in a Neubauer chamber.

### Drop-seq procedure, single-cell library generation and sequencing

For a single-cell encapsulation in NADIA instrument (Dolomite Bio, #3200590), we loaded 75.000 cells in a volume of 250 µl (300.000 cells/ml) and 150.000 Macosko oligodT beads (ChemGenes Corporation, #Macosko-2011-10(V+)) in 250 µl (600 beads/µl) previously washed and resuspended in lysis buffer (6% w/v Ficoll PM-400, 0.2% v/v Sarkosyl, 0.02 M EDTA, 0.2 M Tris pH 7.5 and 0.05 M DTT in nuclease-free water). Cells and beads co-flowed in the microfluidic chip of the device with a capture efficiency of 5-7%.

Immediately after the droplet emulsion breakage, the RNAs captured by the oligodT are reverse transcribed (maxima H RT Master Mix, Thermo, #EP0751). Collected single-cell transcriptomes attached to microparticles (STAMPS) were then splitted in pools of 4000 and amplified for 9 or 11 PCR cycles depending on the bead batch used for the encapsulation (01 and 02 respectively). After cDNA purification with AMPure XP Beads (Agencourt, #A63881), quantification with Qubit dsDNA HS Assay (Thermo, #Q32851) and fragment size check-up using a 4200 TapeStation System (Agilent, #G2991BA), Nextera XT DNA Library Prep Kit (Illumina, #FC-131-1096) was used for the tagmentation of 600 pg of cDNA, illumina adaptor tagging and amplification. The size of Nextera libraries after being purified with AMPure XP Beads was determined using a 4200 TapeStation System and quantified with Qubit dsDNA HS Assay.1.8 pM of pooled libraries was sequenced on Illumina NextSeq 550 sequencer using Nextseq 550 High Output v2 kit (75 cycles) (Illumina, #20024906) in paired-end mode; 20 bp for Read 1 using the custom primer Read1CustSeqB^34^ (cell barcode and UMI) and 64 bp for Read 2, and 8 bp for i7 index.

### scRNA-seq data pre-processing

scRNA-seq libraries were processed using *Drop-seq_tools* 2.3 pipeline^22^ to generate Digital Gene Expression (DGE) matrices. First, *Drop-seq tools* were used to generate the index and the annotation files for the hg38 assembly version of the human genome using Ensembl version 100 annotation^35^ as reference. Next, fastq files containing paired-end reads were merged into a single unaligned BAM file using picard tools v 2.18.14^36^. Using the Drop-seq toolkit with default parameters, reads were then tagged with the cell and the molecular barcodes, trimmed at the 5’ end to remove adapter sequences and at the 3’ end to remove polyA tails. Next, reads were mapped to the human genome (version hg38) with *STAR* version 2.7.0.a^37^. Resulting bam files were tagged with the annotation metadata files to identify reads overlapping genes. Finally, cell barcode correction was done using the programs *DetectBeadSubstitutionError* and *DetectBeadSynthesisErrors* also with default parameters. To estimate the number of cells obtained during the single-cell encapsulation, we used a “knee plot” using as input the number of uniquely mapped reads assigned to the top *N* barcodes, where *N* is at least 5 times the number of expected cells. The estimated number of cells obtained with this procedure was then used to generate a DGE. Two DGE matrices were generated for each dataset, one containing all UMIs overlapping genes using the parameters *LOCUS_FUNCTION_LIST=INTRONIC LOCUS_FUNCTION_LIST=INTERGENIC* and another one containing all UMIs overlapping introns using the parameters L*OCUS_FUNCTION_LIST=null LOCUS_FUNCTION_LIST=INTRONIC*.

### Filtering of low-quality cells and doublets

DGE expression matrices were analyzed using Seurat v 4.0.2^38^. First, we generated Seurat objects for each dataset and used DoubletFinder^39^ to remove doublets. The parameters and doublets identified in each dataset are detailed in table S3. After doublet removal, we merged individual objects to perform a joint analysis. Initially, all genes expressed in less than 3 cells were removed. After manual inspection, all cells with less than 200 detected genes, more than 17000 UMIs, a percentage of mitochondrial transcripts > 7.5%, or a ribosomal content > 40% were discarded. Additionally, we fitted a linear model to describe the relationship between the log number of UMIs and the log number of genes detected per cell. All cells with a residual smaller than -0.5 (2 cells) were discarded. The final Seurat object obtained contained 14588 cells and 23719 genes.

### Identification of cell populations

We used *Seurat* function to regress out the percentage of mitochondrial transcripts, the number of genes, the number of UMIs and the preservation method. To normalize data we used the *LogNormalize* method and multiplied by a scale factor of 10000. We then selected the 2000 most variable genes to calculate 100 principal components (PCs). We used the *ElbowPlot* function to manually inspect the amount of variability explained by each PC and select the first 20 PCs that were used to build the kNN graph and compute the UMAP plot using 500 training epochs (iterations). To eliminate the batch effects affecting the identification of shared cell populations across datasets we used Harmony package ^27^. The function *RunHarmony* was applied on the filtered and processed object, providing the samples as the variable to integrate. By inspecting the updated Elbow plot, we selected the first 20 corrected PCs to perform the clustering. We used the package clustree^40^ to inspect the clustering results at different resolutions from 0.1 to 1 and chose a final resolution of resolution 0.8 where we obtained 13 cell populations. To calculate the top markers for each cluster, we used *FindAllMarkers* function from Seurat with only positive markers and the rest default parameters. Statistically significant markers with adjusted p-values of less than 0.05 were selected.

### Stress and apoptosis gene signatures

We used the function *AddModuleScore* with default parameters from Seurat package ^38^ to assess if the different fixation methods induced stress or favored apoptosis among the cells. This function compares the expression of a set of given genes with random sets of genes with similar expression in the dataset to calculate an enrichment. For the apoptosis signature, we built a gene signature including the genes BCL2, TNF, TP53, CASP3, BAX, CASP8, FAS^41^. For the stress signature, we used the following immediate early genes FOS, JUN, EGR1, UBC, HSPA1B, BTG2, IER2, ID3^28^.

### Cell composition analysis

To assess changes in cell composition we used scCODA^5^. We chose cluster 7 as a reference cell type as this cluster had a good number of cells and a very low amount of dispersion (expressed as differences between groups). To ensure the results were consistent and reproducible we ran scCODA^5^ 10 times using the Hamiltonian Monte Carlo (HMC) sampling method and we averaged the results.

### Correlation analysis

To assess the correlation across datasets, we compared the expression profiles of all cells from each dataset globally and at the cluster level. For that purpose, we computed pseudo bulk expression values for all genes by adding the counts from all the cells within a specific cluster/dataset. Afterwards, we log transformed the counts *c* using a pseudocount so that the normalized expression *n* was *n = log (c + 1)*. We used these normalized expression values to calculate the Pearson correlation coefficient per cluster/sample using *cor* function in R.

### Measuring the capture efficiency of bead batches

Different batches of beads can have different mRNA capture efficiency. In order to evaluate the impact of using two different batches in scRNA-seq encapsulations, we measured the capture efficiency of both bead batches. We encapsulated the same sample twice using both bead batches used in the paper. After RT-PCR, we amplified 4000 STAMPS by PCR using 9, 10, 11 or 12 cycles independently. After AMPure XP Beads purification, the cDNA of each PCR was quantified by Qubit dsDNA HS Assay. Our results demonstrate that to obtain a similar cDNA concentration with the two bead batches, we needed to increase by 2 the number of PCR when using batch 02 (Figure S2).

## Data availability

All raw and processed scRNA-seq data generated for this study can be found in GEO database under accession number GSE209947.

## Author contributions

AGF and MP designed the project and planned experiments. AGF performed all experimental work. MH and MP performed computational data. LM performed compositional analysis of the samples. MP acquired funding, supervised, and coordinated the work. AGF and MP interpreted the results. MP wrote the manuscript with input from all authors.

## Acknowledgements

We thank all members from the Plass Lab for useful comments and critical discussion. We also thank Yvonne Richaud-Patin from the Regenerative Medicine Program at IDIBELL and Dr. Zomeño from IDIBELL’s Advanced Cell and Tissue Culture platform for their help with iPSC cell culture. We thank Dr. Iglesias, Dr. Kim and Dr. Sebé-Pedrós for their support in setting up ACME and vivoPHIX protocols for cell preservation. This research was funded by a research project from the State R&D Program Research Challenges from the Spanish Ministry of Science, Innovation and Universities (Grant number: PID2019-108580RA-I00). MP work is supported by a Ramón y Cajal contract of the Spanish Ministry of Science and Innovation (RYC2018-024564-I). We thank CERCA Program / Generalitat de Catalunya for IDIBELL institutional support.

## Supplementary material

**Figure S1.**
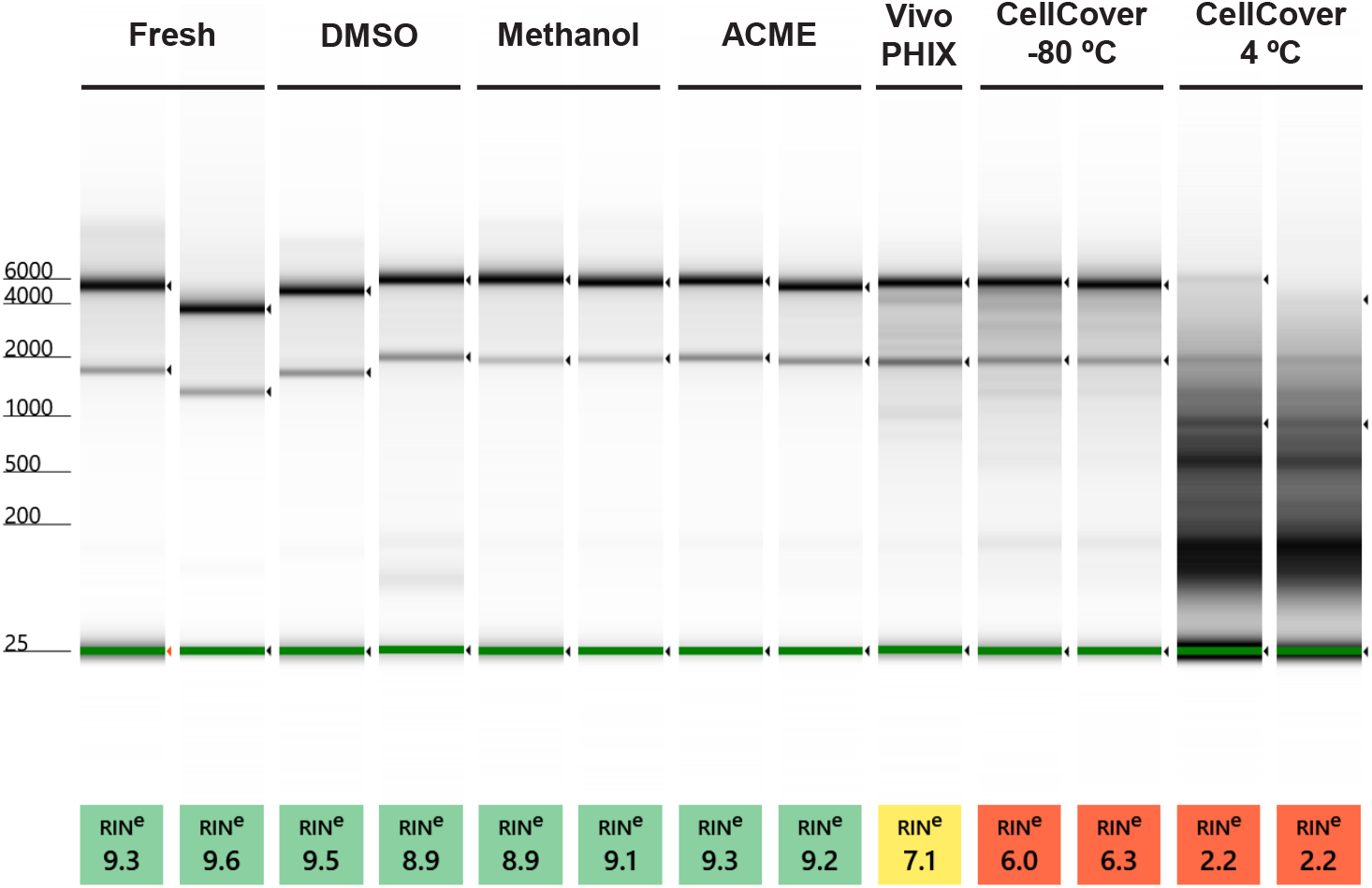
RNA quality control of preserved samples. Gel image of the RNA profile of different samples obtained from a High Sensitivity D5000 ScreenTape assay using a 4200 Agilent Tape Station system. Bands marked with < represent 28S, 18S and the lower marker. The RNA quality (RIN values) for each sample is shown below the gel. The RNA quality of DMSO, methanol and ACME samples is very high and equivalent to that of Fresh samples. vivoPHIX shows a medium RNA quality while CellCover samples at -80ºC or 4ºC show lower RNA quality, specially at 4ºC.

**Figure S2.**
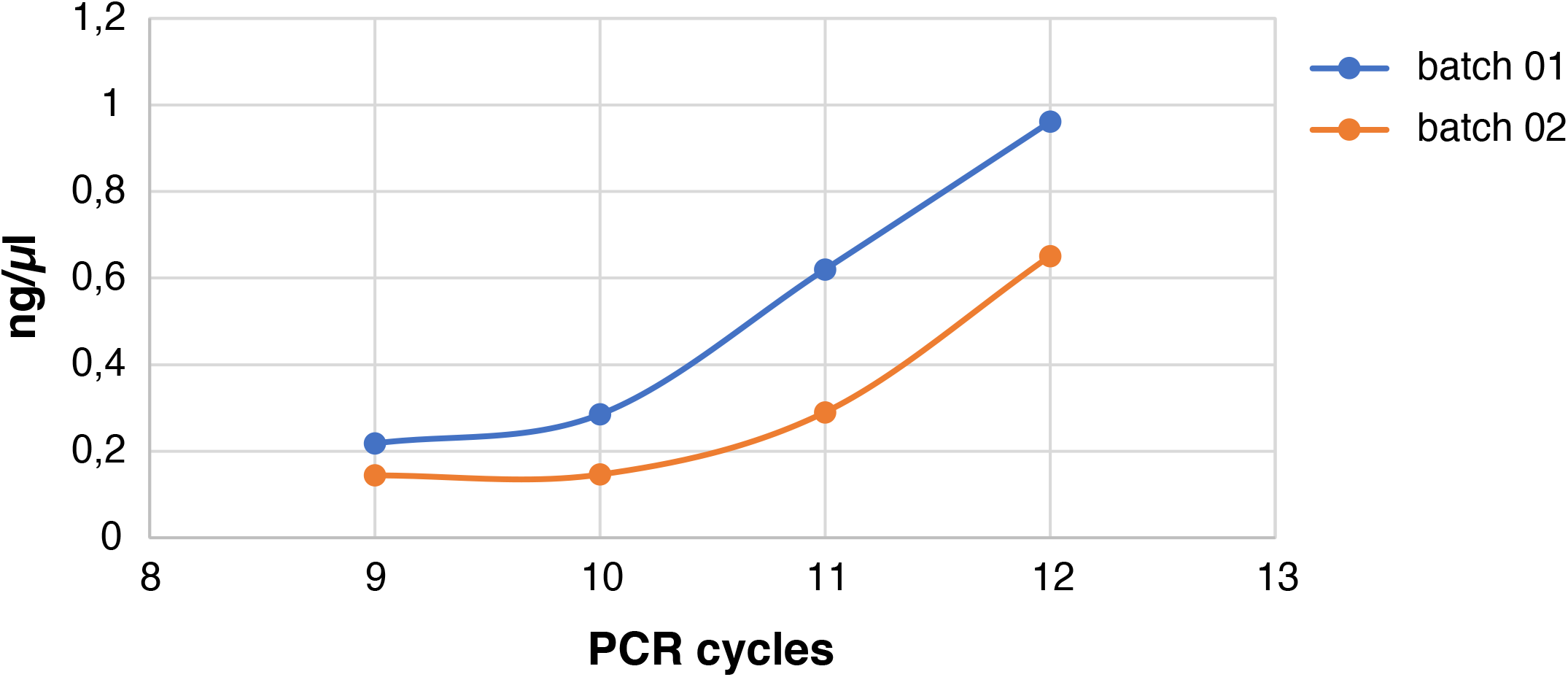
Comparison of the capture efficiency of the 2 bead batches used in the project. The plots show the concentration of cDNA in ng/µl as a function of PCR cycles. Beads from batch 01 have a higher capture efficiency than batch 02, which is reflected in a higher amount of genes and UMIs in the samples processed with these beads (Table S1).

**Figure S3.**
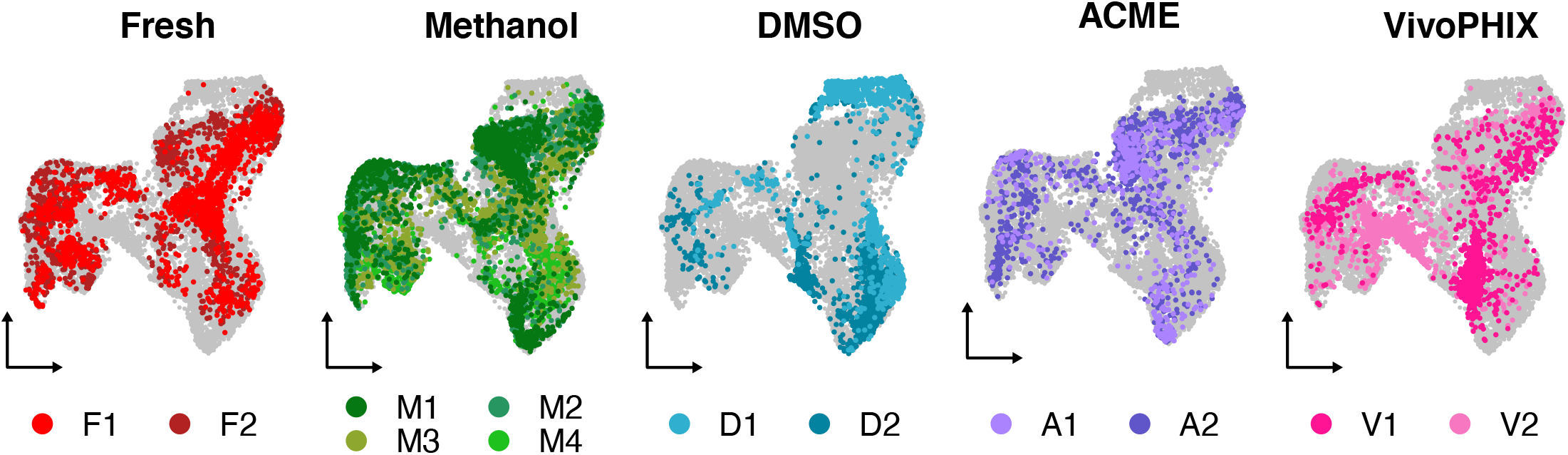
Cell preservation induces biases in gene expression that affect sample comparison. UMAP plots showing the distribution of cells on the UMAP for each of the preservation methods used. Each plot highlights the distribution of cells from individual replicates obtained using the same preservation method.

**Figure S4.**
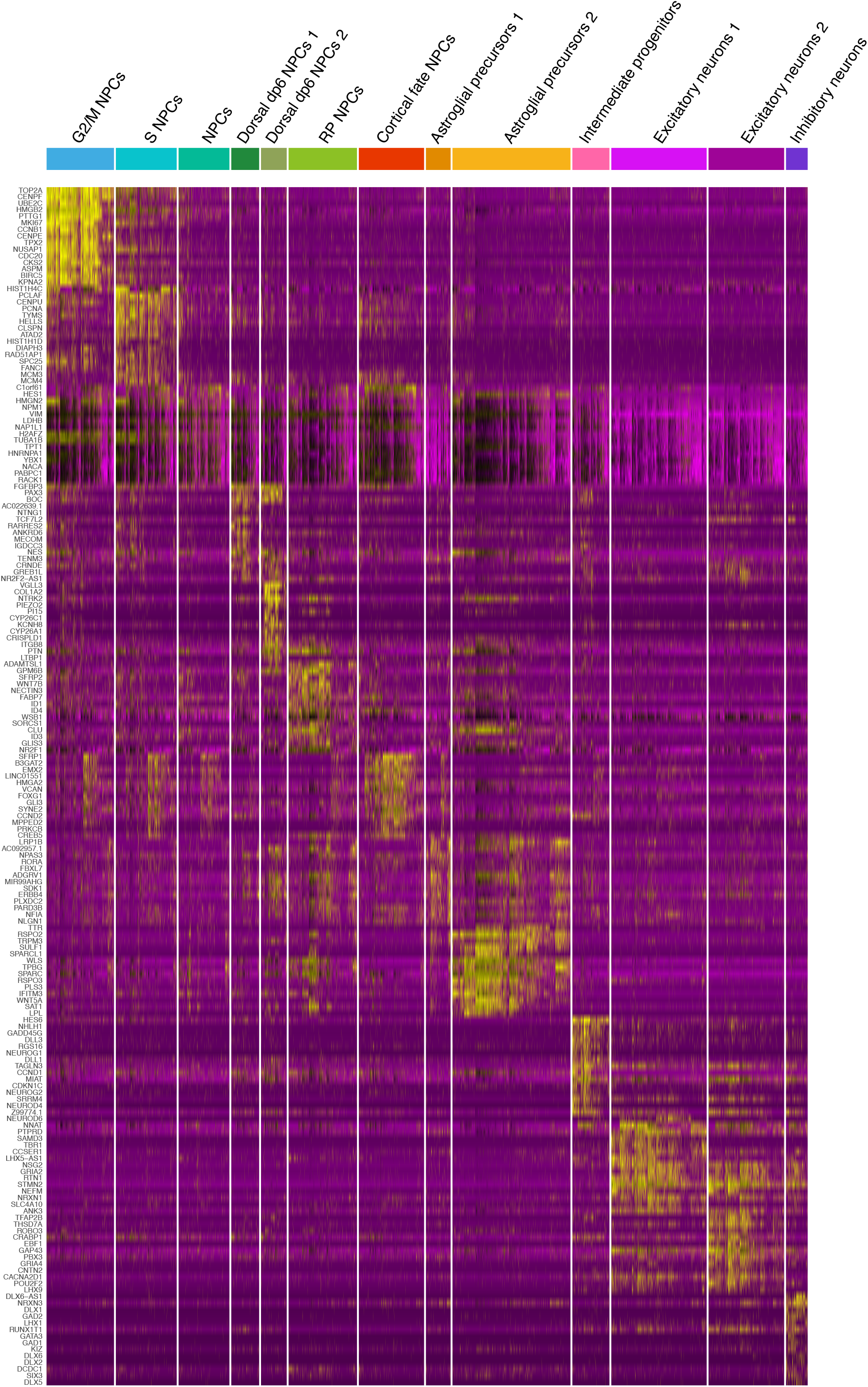
Top 15 markers identified for each cluster. Heatmap plot showing the expression of the top 15 markers identified in each cluster in each cell. Expression values range from purple (low expression) to yellow (high expression).

**Figure S5.**
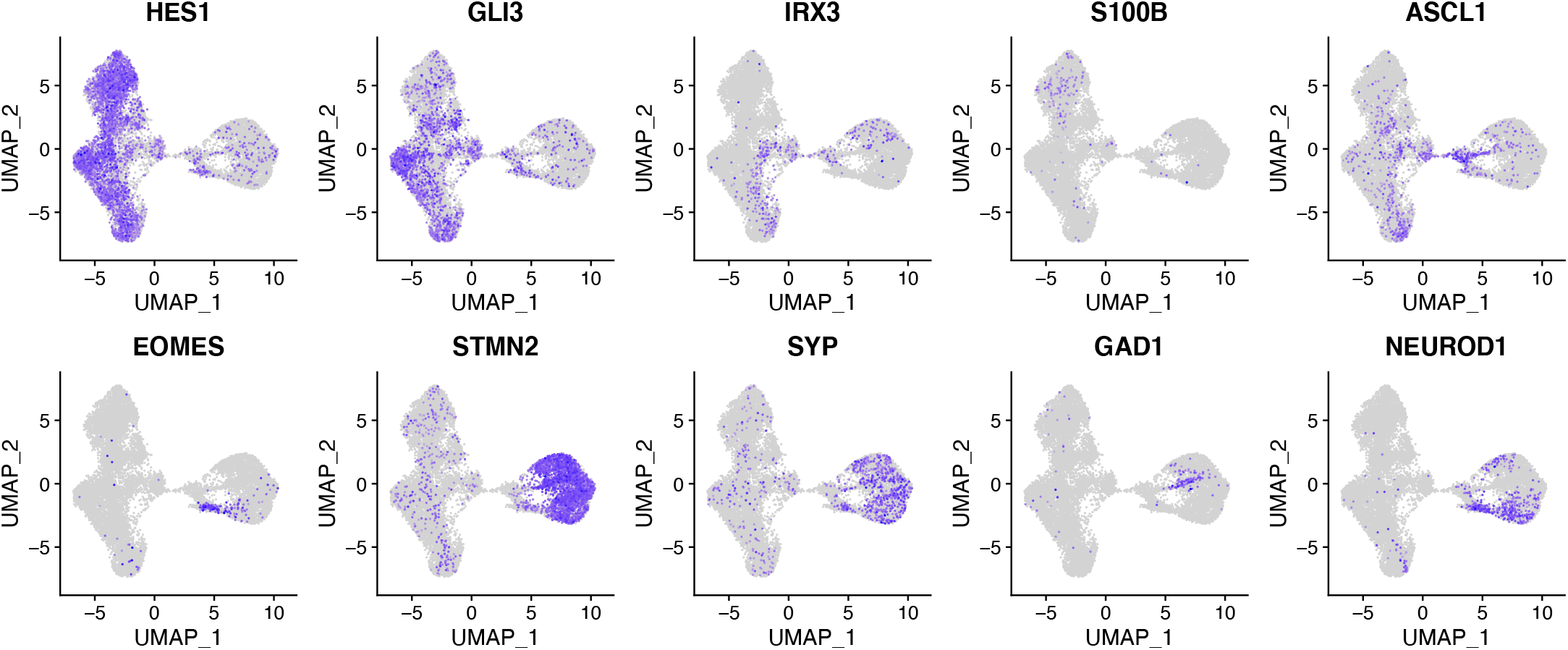
Additional feature plots validating the identity of the identified clusters. Feature plots of known markers that have been used to identify the cell populations obtained (Figure3a). These markers validate the identity of NPCs (HES1), cortical fate NPCS (GLI3), dorsal fate NPCs (IRX3), astrocytes (S100B), intermediate progenitor neurons (ASCL1, EOMES), immature (STMN2) and mature (SYP) neurons, and inhibitory (GAD1) and excitatory neurons (NEUROD1).

**Figure S6.**
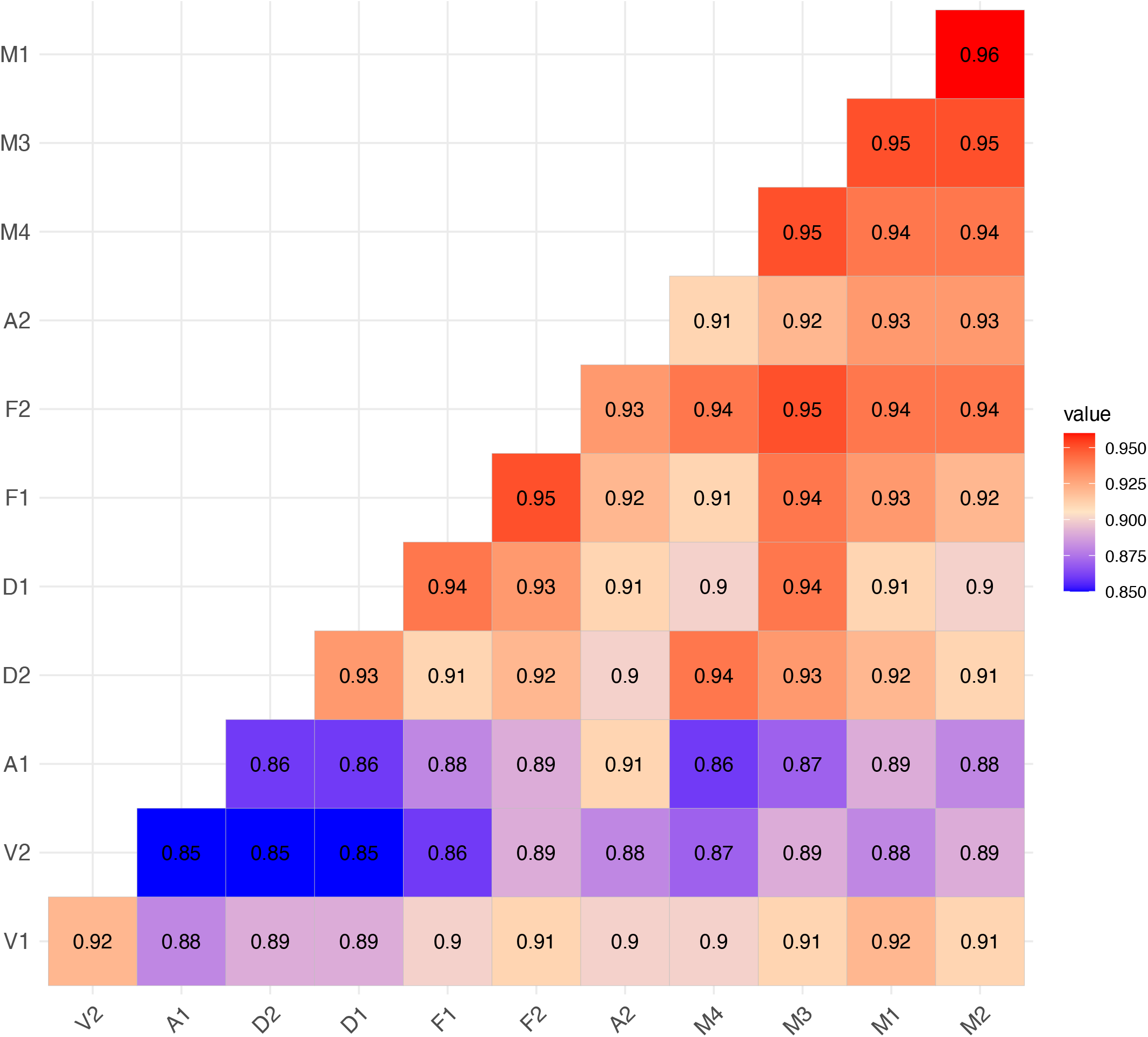
Gene expression correlation across samples. Heatmat plot showing the pseudo-bulk gene expression correlation across samples. Inside each cell, the Pearson correlation coefficient is shown. Both ACME and vivoPHIX samples show lower correlation with all the other samples. Yet, the average correlation across all samples is quite high (r >= 0.85).

**Table S2.** Estimated amount of UMI and Genes at 7500 reads per cell.

| Name | Estimated Genes | Estimated UMIs |
| --- | --- | --- |
| F1 | 886 | 1414 |
| F2 | 728 | 1097 |
| M1 | 840 | 1254 |
| M2 | 855 | 1292 |
| M3 | 1095 | 1798 |
| M4 | 1205 | 2162 |
| D1 | 1325 | 2375 |
| D2 | 1471 | 3021 |
| A1 | 267 | 379 |
| A2 | 425 | 674 |
| V1 | 505 | 690 |
| V2 | 225 | 280 |

**Table S3.** Parameters and number of doublets identified using DoubletFinder.

| Sample | PCs | pK | No. of doublets |
| --- | --- | --- | --- |
| F1 | 11 | 0.01 | 132 |
| F2 | 10 | 0.04 | 100 |
| M1 | 14 | 0.01 | 138 |
| M2 | 10 | 0.02 | 131 |
| M3 | 11 | 0.3 | 137 |
| M4 | 7 | 0.02 | 98 |
| D1 | 12 | 0.1 | 93 |
| D2 | 8 | 0.22 | 101 |
| A1 | 6 | 0.3 | 98 |
| A2 | 9 | 0.01 | 114 |
| V1 | 12 | 0.3 | 118 |
| V2 | 11 | 0.005 | 229 |

**Table S5.** Gene markers used to identify cell populations in the scRNA-seq.

| GENE NAME | CELL TYPE | REFERENCES |
| --- | --- | --- |
| **SOX2** | Neural precursor | 42 |
| **HES1** | Neural precursor | 43 |
| **TOP2A** | Proliferation | 44 |
| **PCNA** | Proliferation | 44 |
| **FOXG1** | Cortical fate | 45 |
| **GLI3** | Cortical fate | 46 |
| **PAX3** | Dorsal fate | 47 |
| **IRX3** | Dorsal fate | 47 |
| **PTN** | Roof plate | 47 |
| **ADGRV1** | Astroglia | 45 |
| **GJA1** | Astroglia | 45 |
| **S100B** | Astrocytes | 45 |
| **HES6** | Intermediate progenitor | 45 |
| **ASCL1** | Intermediate progenitor | 48 |
| **EOMES** | Intermediate progenitor | 48 |
| **DCX** | Immature neurons | 48 |
| **STMN2** | Immature neurons | 1 |
| **SYP** | Mature neurons | 1 |
| **GAD1** | Inhibitory neurons | 1 |
| **GAD2** | Inhibitory neurons | 1 |
| **SLC17A6** | Excitatory neurons | 48 |
| **NEUROD1** | Excitatory neurons | 48 |

